# Combined analysis of genome sequencing and RNA-motifs reveals novel damaging non-coding mutations in human tumors

**DOI:** 10.1101/200188

**Authors:** Babita Singh, Juan L. Trincado, PJ Tatlow, Stephen R. Piccolo, Eduardo Eyras

## Abstract

A major challenge in cancer research is to determine the biological and clinical significance of somatic mutations in non-coding regions. This has been studied in terms of recurrence, functional impact, and association to individual regulatory sites, but the combinatorial contribution of mutations to common RNA regulatory motifs has not been explored. We developed a new method, MIRA, to perform the first comprehensive study of significantly mutated regions (SMRs) affecting binding sites for RNA-binding proteins (RBPs) in cancer. Extracting signals related to RNA-related selection processes and using RNA sequencing data from the same samples we identified alterations in RNA expression and splicing linked to mutations on RBP binding sites. We found SRSF10 and MBNL1 motifs in introns, HNRPLL motifs at 5’ UTRs, as well as 5’ and 3’ splice-site motifs, among others, with specific mutational patterns that disrupt the motif and impact RNA processing. MIRA facilitates the integrative analysis of multiple genome sites that operate collectively through common RBPs and can aid in the interpretation of non-coding variants in cancer. MIRA is available at https://github.com/comprna/mira.

## Introduction

Cancer arises from genetic and epigenetic alterations that interfere with essential mechanisms of the normal life cycle of cells such as DNA repair, replication control, and cell death (1). The search for driver mutations, which confer a selective advantage to cancer cells, is generally performed in terms of the impact on protein sequences (2). However, systematic studies of cancer genomes have highlighted mutational processes outside of protein-coding regions (3–5) and tumorigenic mutations at non-coding regions have been described, like those in the TERT promoter (6,7). However, a major challenge remains to more accurately and comprehensively determine the significance and potential pathogenic involvement of somatic variants in regions that do not code for proteins (8). Current methods to detect driver mutations in non-coding regions are based on 1) the recurrence of mutations in predefined regions in combination with measurement of potential functional impacts (4,9–11), 2) recurrence in combination with sequence conservation or polymorphism data (12,13), or 3) the enrichment of mutations with respect to specific mutational backgrounds (5,14,15); and some of them combine such approaches (5). However, these methods have so far been restricted to individual positions rather than combining the contributions from multiple functionally equivalent regulatory sites. Additionally, the impact on RNA processing measured from the same samples have not been evaluated

RNA molecules are bound by multiple RNA binding proteins (RBPs) with specific roles in RNA processing, including splicing, stability, localization and translation, which are critical for the proper control of gene expression (16). Multiple experimental approaches have established that RBPs generally interact with RNAs through short motifs of 4-7 nucleotides (17–19). These motifs occur anywhere along the precursor RNA molecule (pre-mRNA), including introns, protein coding regions, untranslated 5’ and 3’ regions, as well as in short and long non-coding RNAs (20,21). Mutations on RNA regulatory sequences can impact RNA processing and lead to disease (22). However, studies carried out so far have mainly focused on sequences around splice sites (23), or in protein-coding regions (24). *In vitro* screenings of sequence variants in exons has revealed that more than 50% of nucleotide substitutions can induce splicing changes (25,26), with similar effects for synonymous and non-synonymous sites (26). Since RBP binding motifs are widespread along gene loci, and somatic mutations may occur anywhere along the genome, it is possible that mutations in other genic regions could impact RNA processing and contribute to the tumor phenotype. Mutations and expression alterations in RBP genes have an impact on specific cellular programs in cancer, but it is not known whether mutations in RBP binding sites along gene loci are frequent in cancer, could damage RNA processing, and contribute to oncogenic mechanisms.

To understand the effects of somatic mutations on RNA processing in cancer at a global level we have developed a new method, MIRA, to carry out a comprehensive study of somatic mutation patterns along genes that operate collectively through interacting with common RBPs. Compared with other existing approaches to detect relevant mutations in non-coding regions, our study provides several novelties and advantages: 1) we searched exhaustively along gene loci, hence increasing the potential to uncover deep intronic pathological mutations; 2) we studied the enrichment of a large compendium of potential RNA regulatory motifs, allowing us to identify potentially novel mechanisms affecting RNA processing in cancer; 3) we showed that multiple mutated genomic loci potentially interact with common RBPs, suggesting novel cancer-related selection mechanisms; and 4) unlike previous methods, we used RNA sequencing data from the same samples to measure the impact on RNA processing. Our study uncovered multiple mutated sites associated to common RBPs that impact RNA processing with potential implications in cancer, and revealed a new layer of insight to aid in the interpretation of non-coding variants in cancer genomes.

## Results

### Unbiased search for significantly mutated regions (SMRs) along gene loci

We performed an exhaustive detection of mutation enrichment using overlapping genomic windows of 7 nucleotides (7-mer windows) along each gene locus (Fig. 1a) (Figure S1) (Methods). Using a dataset of somatic mutations from whole-genome sequencing (WGS) from 505 samples for 14 tumor types (10) (PAN505) (Table S1), we performed a double statistical test. First, to account for local variations in mutational processes we compared each 7-mer window against the mutation rate in the entire locus, and selected those with p-value < 0.05 after correcting for multiple tests (Figure S2a) (Methods). Secondly, to account for nucleotide biases, we compared the mutation count in each 7-mer window with the expected count calculated from the mutation rate per nucleotide to define a nucleotide bias (NB) score per window (Methods). Of the 140,704 windows with 3 or more mutations, 93,497 (66%) showed NB-score > 6, whereas of the 45,916,437 windows with 1 mutation, which we considered to reflect the background, 1,557,310 (3%) had NB-score > 6 (Fisher’s exact test p-value = 0, odds-ratio = 369.5) (Figure S2b). Using the filters of p-value < 0.05 and NB score > 6, our exhaustive search produced 78,352 significant 7-mer windows in 8,159 genes. We further separated 7-mer windows according to whether they were in a coding sequence (CDS), a 5’ or 3’ untranslated region (5UTR/3UTR), an exon in a long non-coding RNA (EXON), an intron (INTRON), or overlapping a 5’ or 3’ splice site (5SS/3SS) and clustered them into significantly mutated regions (SMRs) (Fig. 1a), producing a total of 20,307 SMRs, harboring a total of 41,756 substitutions (Figure S3) (Tables S2 and S3). Most of the predicted SMRs were 7-15 nucleotides long (Figure S4), and the majority of SMRs were in introns and in exons of non-coding RNAs (EXON) (Table 1).

**Table 1.** Significant mutated regions (SMRs). For each region type, we indicate the number of SMRs predicted (Total), SMRs with stranded enriched motifs (with motifs), with stranded enriched motifs that we could label (with labeled motifs), and with a significant association to an RNA-processing change (impact on RNA). This latter case includes changes in exon-exon junctions and transcript expression.

| SMRs | Total | With motifs | With labeled motifs | Impact on RNA |
| --- | --- | --- | --- | --- |
| 3SS | 823 | 341 | 335 | 2 |
| 3UTR | 294 | 44 | 44 | 7 |
| 5SS | 1054 | 546 | 521 | 11 |
| 5UTR | 119 | 45 | 45 | 5 |
| CDS | 208 | 10 | 10 | 3 |
| EXON | 335 | 119 | 114 | 12 |
| INTRON | 17474 | 3176 | 1812 | 427 |

**Figure 1.**
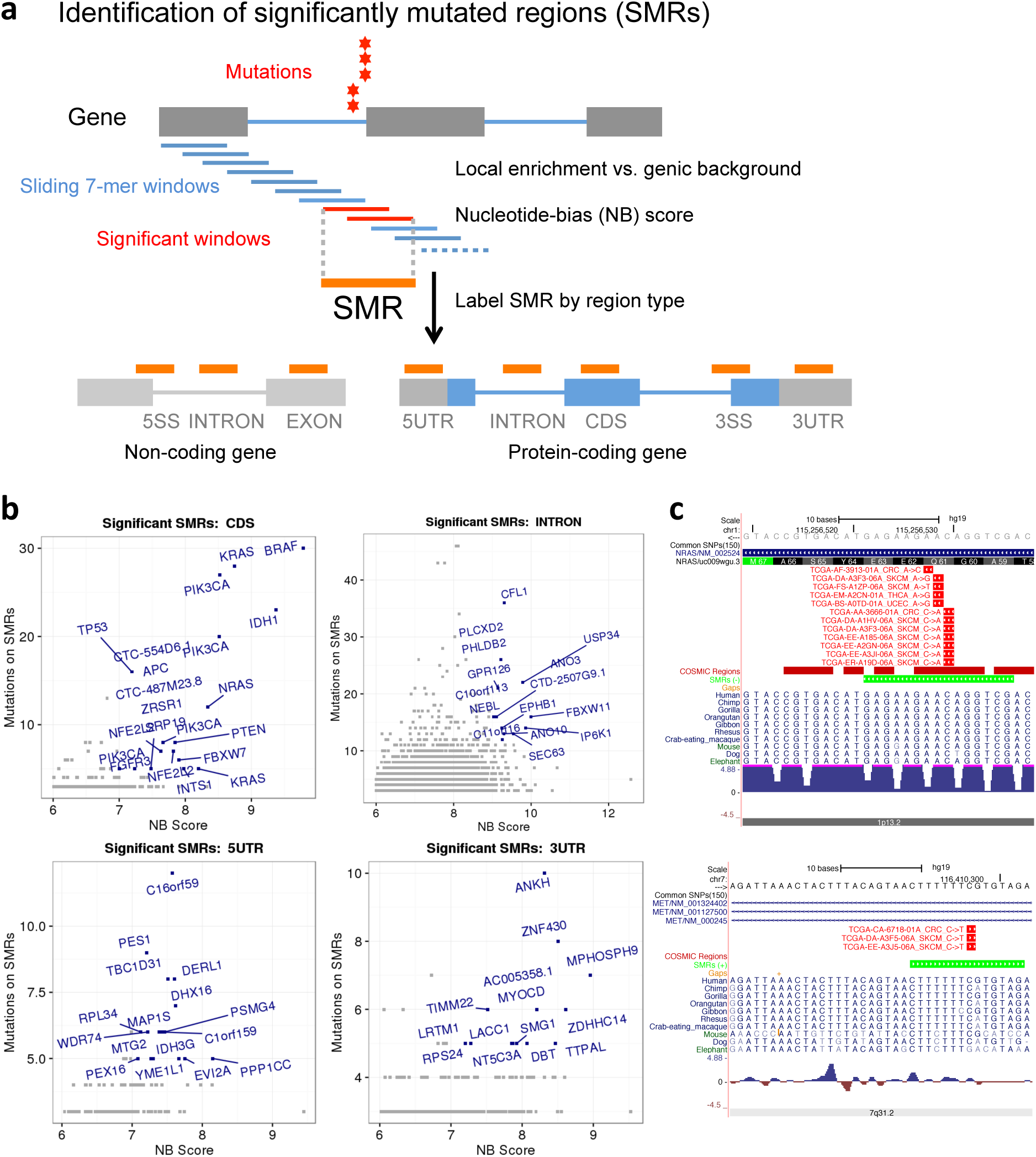
Systematic identification of RNA-related significantly mutated regions (SMRs). **(a)** Short k-mer windows (k=7 in our study) along genes are tested for the enrichment in mutations with respect to the gene mutation rate and the local nucleotide biases. Significant windows are clustered by region type into significantly mutated regions (SMRs). For each SMR we give the NB-score (x axis) and the number of mutations (y axis), (b) We show the SMRs detected in CDS regions, introns (INTRON), 5’ UTRs (5UTR) and 3’UTRs (3UTR). All SMRs detected are shown, but we only show the gene name for the SMRs with nucleotide-bias (NB) score > 6 and with 5 or more mutations, except for the INTRON SMRs, where we highlight the cases with 12 or more mutations. **(c)** Examples of a CDS SMR in *NRAS* and an INTRON SMR in *MET*. The UCSC screenshots show the SMR (green) and the mutations detected (red).

We tested our predicted SMRs for possible biases. We calculated for each SMR the DNA replication timing, which is known to correlate with somatic mutations in cancers and can be a source of artifacts (27,28), and observed no association with mutation count (Figure S5). Another potential source of artifacts is the relation between gene expression and mutation rates (27). We used RNA sequencing (RNA-seq) data from the same samples to measure the expression of SMR-containing transcripts and observed no association between the mutation count and expression (Figure S6). We further used LARVA (14) to test our SMRs using a statistical model that accounts for over-dispersion of the mutation rate and replication timing (Methods). We observed an overall high similarity between the significance provided by MIRA and LARVA (Figure S7). In particular, we found a strong agreement for INTRON SMRs (Pearson’s R = 0.83). Additionally, we analyzed our SMRs with OncodriveFML (11). Most of the CDS and EXON SMRs were identified as significant by OncodriveFML, whereas not as many INTRON, 5UTR or 3UTR SMRs were seen as significant (Figure S8) (Table S2), suggesting that mutations falling on INTRON or UTR regions may impact function in a different way that has not been considered yet.

### Predicted SMRs recovered known and novel mutational hotspots

We found SMRs in 501 cancer-driver genes out of 889 collected from the literature (29) (Fig. 1b) (Figure S9), which is more than expected by chance (Fisher’s exact test p-value = 7.24e-109, odds-ratio = 4.67) compared to the 8870 non-cancer genes (out of 40973) that do not harbor SMRs. We observed CDS SMRs in 34 cancer genes identified previously (11,13), including *BRAF*, *IDH1*, *KRAS*, *PIK3CA*, *SF3B1*, *CTNNB1*, *TP53* and *KRAS* (Fig. 1b), as well as in in cancer genes not found previously, including *NRAS*, *EP300* and *ATM* (Fig. 1c). From the 133 genes with predicted 5UTR SMRs, 17 were identified previously (11) and 8 corresponded to cancer genes, including *SPOP* and *EEF1A1*. We also found 3UTR SMRs in 28 cancer genes, including *CTNNB1* and *FOXP1*. From the 519 SMRs in EXON SMRs, 18 were located in cancer gene loci. In 42 lncRNAs related to cancer (15), we found only one EXON SMR in *TCL6*, but 11 INTRON SMRs, suggesting that lncRNAs introns could be more relevant than previously anticipated. As our analysis was exhaustive along the entire gene loci, we recovered many more INTRON SMRs than in previous reports, 317 of which within cancer genes, including *NUMB, ALK*, *EPHB1*, *ARID1A* and *MET* (Fig. 1c). Additionally, 5 of the previously reported intronic mutations (11), we classified as 5SS/3SS SMRs, with *ATG4B*, *NF1* and *TP53* having both types of SMRs.Finally, we found 62 5SS SMRs and 53 3SS SMRs in cancer genes, including *MET, CHEK2*, *BRCA1*, *VEGFA*, *RB1*, *CDKN2A*, as well as *TP53*, *PTEN* and *CHD1*, which were described before to have cancer mutations at splice-sites (23). In summary, our SMRs provide a rich resource with potential to uncover new relevant non-coding alterations in cancer.

### Somatic mutations occur frequently on RBP binding motifs

To further understand the properties of our SMRs we calculated the mutation frequencies at tri-nucleotides considering the strand of the gene in which the SMR was defined. We observed an enrichment of C>T and G>A mutations on SMRs (Fig. 2a, upper panel), which was recapitulated in the reverse-complement triplets, indicating a strong contribution from DNA-related selection processes. To identify SMRs that reflect RNA-related selection processes we studied sequence motifs potentially related to RNA processing. We performed an unbiased k-mer enrichment analysis in SMRs with k=6 (Methods). Further, to select those associated with RNA rather than DNA, we reverse-complemented all SMRs and control sequences and repeated the enrichment analysis, and 6-mers enriched in both calculations for the same region-type were discarded (Figure S1c). We found a total of 357 enriched 6-mers (Table S4) in 3546 SMRs (Table S5). These enriched 6-mers showed a different mutational pattern compared to all SMRs (Fig. 2a) (Table S6). The symmetry between triplets and their reverse complements was no longer present, and there was an enrichment of mutations at AGA and TCC triples, probably reflecting RNA-related selection processes.

**Figure 2.**
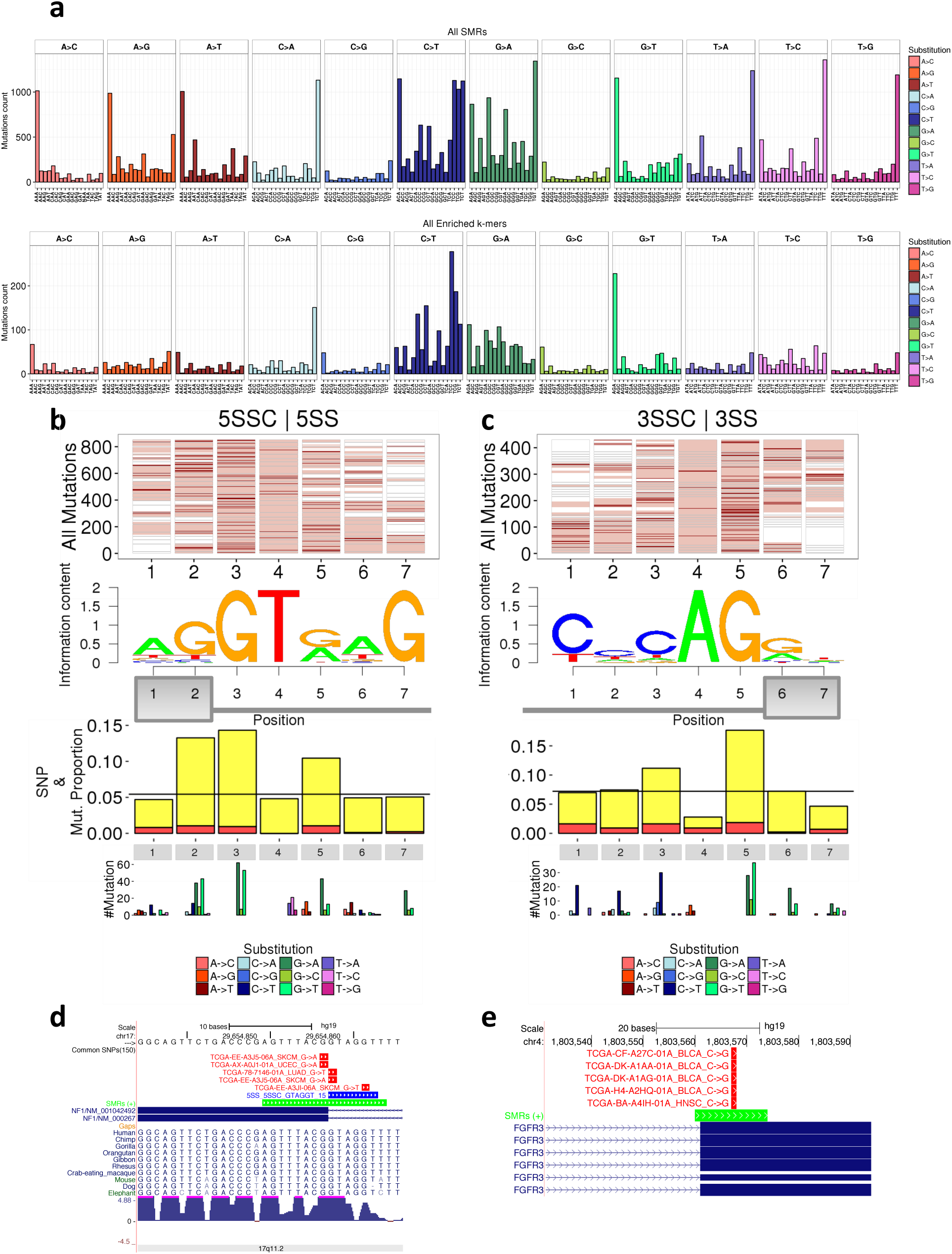
Enriched splice-site motifs in significantly mutated regions (SMRs). **(a)** Upper panel: mutation pattern in SMRs. For each nucleotide substitution we give the total count of substitutions observed in SMRs separated according to the nucleotide triplet in which it occurs. SMRs are stranded; hence we give the substitutions according to the SMR strand. Lower panel: mutation patterns in enriched 6-mers in SMRs. For each nucleotide substitution we give the total count observed separated according to the nucleotide triplet in which it occurs. Since the 6-mers are stranded, give the substitutions according to the 6-mer strand. **(b)** We show the logos for the splice site motifs significantly enriched in 5SS and 3SS SMRs, i.e. they appear more frequently in 5SS or 3SS SMRs than in the corresponding controls. The barplots below show the proportion of all somatic mutations in the 6-mers (y axis) that fall on each position along the motif logo. In orange indicate those somatic mutations that coincide with a germline SNP. **(c)** The plots show for each position the number of splice-sites with each type of substitution indicated with a color code below. **(d)** We give two examples of mutations found at splice sites motifs in the genes *NF1* and *FGFR3*. Above the gene track we show the significantly mutated region (SMR) (green track), the enriched motif found in the SMR (blue track), and the somatic mutations (read track). For each mutation we indicate the patient identifier, the tumor type and the substitution.

From the 74 enriched 6-mers found on 5SS SMRs, 46 included the 5’ splice-site consensus GT (5SSC). 5SSC motifs showed a strong conservation of G at the +5 intronic position (position 7 in Fig. 2b), with the highest density of mutations, mostly G>A and G>T, on either side of the exon-intron boundary (Fig. 2b). From the 52 enriched 6-mers found on 3SS SMRs, 36 contained the 3’ splice site (3’ss) consensus AG (3SSC), with strong conservation of C nucleotides at the -5 position of the intron (position 1 in Fig. 2c) and with positions -1 and -3 (3 and 5 in Fig. 2c) being the most frequently mutated positions (Fig. 2c). Among the cases found, there was a 5’ss in *NF1* with mutations in skin (SKCM), lung (LUAD) and uterine (UCEC) tumors (Fig. 2d), and a 3’ss in *FGFR3* with mutations in head and neck (HNSC) and bladder (BLCA) tumors (Fig. 2e). Mutations at splice site motifs where more frequent in lung (LUAD, LUSC) and uterine (UCEC) tumors; 5SSC mutations were also frequent in bladder tumors (BLCA), whereas 3SSC mutations were specifically frequent in colorectal tumors (CRC) (Fig. 3a).

**Figure 3.**
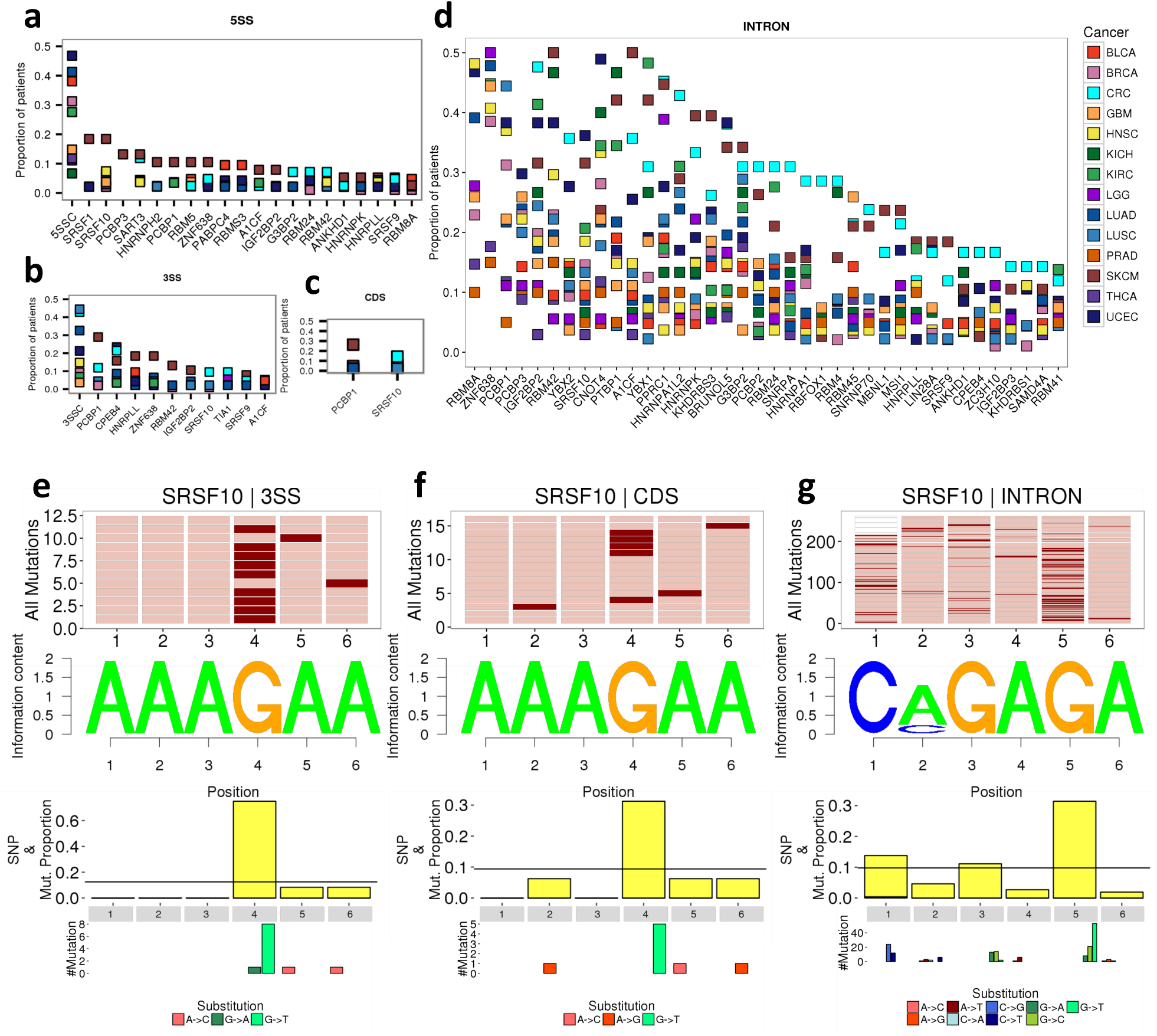
Cancer mutations in enriched RBP motifs. We provide the proportion of samples separated by tumor type (y axis) that have a mutated motif in 5SS **(a)**, 3SS **(b)**, CDS **(c)** and INTRON **(d)** SMRs. In each SMR type we show the enriched motifs. For 5SS and 3SS we indicate the consensus 5’ or 3’ splice site sequences (5SSC/3SSC). The proportions are color coded by tumor type. We show the mutation patterns on SRSF10 motifs in 3SS **(e)**, CDS **(f)** and INTRON **(g)** SMRs. In the upper panel we indicate in dark red the position of the mutations and in light red the positions covered by motif. The barplots below show the proportion of somatic mutations (y axis) that fall on each position along the motif logo. In orange indicate those somatic mutations that coincide with a germline SNP. Below we show for each position the number of motifs with each type of substitution indicated with a color code below.

To identify RBPs that could potentially bind the enriched 6-mers beyond 5SSC/3SSC motifs, we used DeepBind (30) to score the enriched 6-mers using models for 522 proteins containing KH, RRM and C2H2 domains from human, mouse and Drosophila (Methods). We could confidently label 245 (68.6%) of the 357 enriched 6-mers (Table S4). In 5SS and 3SS SMRs, besides the enriched 5SSC/3SSC motifs, we identified binding sites for multiple RBPs (Figs. 3a and 3b); including SRSF10, a splicing factor that regulates an alternative splicing response to DNA damage (31) and PCBP1, which was linked to alternative splicing in cancer (32). SRSF10 and PCBP1 motifs appeared also in CDS and INTRON SMRs (Figs. 3c and 3d) (Figure S11). In 3SS SMRs we also found binding sites for HNRPLL (Fig. 3b), a regulator of T cell activation through alternative splicing (33), which was also found on EXON and INTRON SMRs (Fig. 3d) (Figure S11).

To validate our assignment of RBP labels to SMRs we used data from 91 CLIP experiments for 68 different RBPs (34–40) (Methods). Using the significant CLIP-Seq signals as evidence of protein-RNA interaction, we observed an enrichment of CLIP signal on labeled SMRs compared to SMRs without any RBP assignment (Fisher’s exact test p-value = 4.10e-30, odds-ratio = 1.97). We further tested each specific motif independently for the enrichment of CLIP signals with respect to the other motifs (Table S7) (Figure S10). Motifs associated to splice-site sequences had the highest enrichment of CLIP signal. We also observed that SMRs labeled with CPEB4, SRSF10, PCBP1 or PCBP3 are among the cases with greatest enrichment of CLIP signal. Thus, our labeled SMRs are generally enriched on CLIP-Seq signal compared to non-labeled SMRs, supporting the notion that these SMRs are likely to participate in protein-RNA interactions.

### Somatic mutations show positional biases on RBP binding motifs

We further studied whether particular positions on the identified RBP motifs were more frequently mutated than others. For each RBP, we grouped the SMRs containing the enriched labeled 6-mers and performed a multiple sequence alignment (MSA) to determine the equivalent positions of the motifs across the SMRs. For SRSF10 motifs we found an enrichment of A>G mutations at A positions (Fig. 3d), which was recapitulated at INTRON, CDS and 3SS SMRs (Fig. 3c). On EXON SMRs the most abundant RBP motif was CPEB4, which showed enrichment of T>A mutations, whereas HNRPLL and PCBP1 motifs showed enrichment of C>T mutations (Figure S12). At 5UTR SMRs we found multiple T-and C-rich motifs, predominantly in melanoma (SKCM) (Fig. 4a). In particular, PCBP3 and PTBP1 motifs at 5UTR SMRs, which were characterized by an enrichment of C>T substitutions (Fig. 4b), might be related to the 5’ terminal oligo-pyrimidine tract (5’TOP) motif that is relevant for translational regulation (41,42), and these mutations could indicate an impact on translation. At 5UTR SMRs there were also G-rich motifs (HNRPLL, HNRNPA2B1) with frequent G>A mutations (Figure S13). At 3UTR SMRs (Fig. 4c) we observed frequent CT-rich motifs associated to PTBP1, HNRNPC, PCBP3 and ELAVL1 (HuR) in CRC, BLCA and UCEC patients (Fig. 4d), and AC-rich motifs associated to IGF2BP2 in UCEC and CRC with enrichment of C>T mutations (Figure S14).

**Figure 4.**
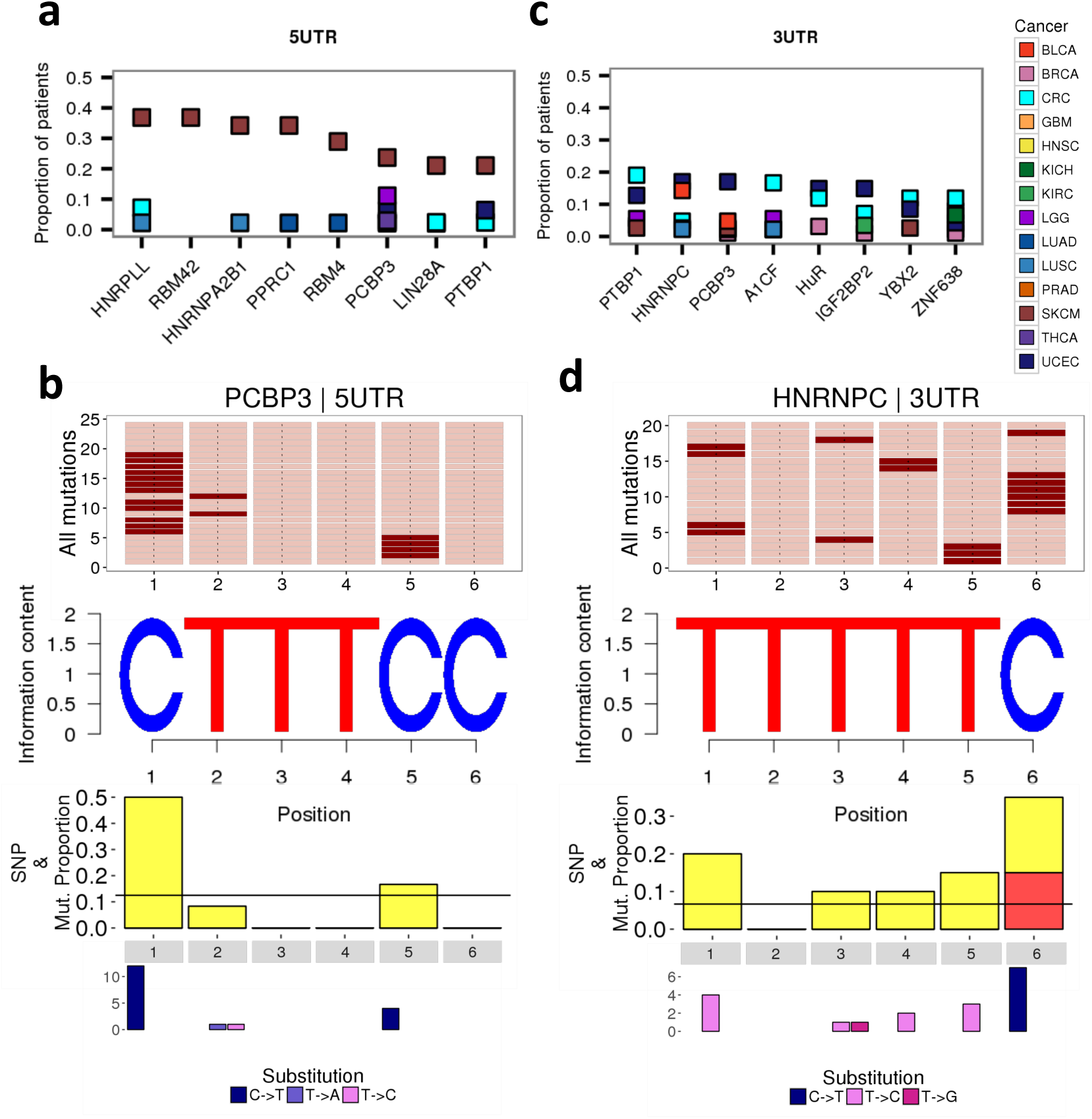
Cancer mutations in enriched RBP motifs in 5UTR and 3UTR SMRs. **(a)** For 5UTR **(a)** and 3UTR **(b)** SMRs we provide the proportion of samples in each tumor type (y axis) that have a mutated RNA binding protein (RBP) motif (x axis). The proportions are color-coded by tumor type. The proportions of SMRs with each RBP motif per tumor type are given in Figure S11. Positional patterns of mutations on **(c)** PCBP3 motifs in 5UTR SMRs, and on **(d)** HNRNPC3 motifs in 3UTR SMRs. In the upper panels we indicate in red the positions covered by the motif and in dark red the position of the mutations. The barplots below show the proportion of somatic mutations (y axis) that fall on each position along the motif logo. In orange we indicate those somatic mutations that coincide with a germline SNP in position (with a different substitution pattern, as the exact matching substitutions were removed). Below we show for each position, the number of motifs with each type of substitution indicated with a color

For each enriched motif, we calculated the enrichment of gene sets (GO Biological Function, Pathways and Oncogenic Pathways) in the genes harboring SMRs with the motif (Table S8). Among the motifs associated with cancer-related functions: 5SSC motifs appeared related to apoptosis and DNA damage response functions, as well as to NFKB activation and PI3K signal cascade (Figure S15). Genes with SMRs harboring SRSF10 motifs showed association to apoptosis and immune response, whereas genes with HNRNPC or HNRNPLL motifs were related to metabolic processes. RBM41-motif containing SMRs are also related to genes involved in metabolic processes, as well as in T-cell activation. These results indicate that cancer mutations on RBP motifs potentially impact a wide range of functions that may lead to specific tumor phenotypes.

### Somatic mutations in RBP binding motifs impact in RNA expression and splicing

To determine the impact of mutations in enriched motifs we tested their association with changes in RNA processing. We first estimated the association of mutations with changes in transcript isoform expression (Methods). From the 20,308 SMRs tested, 148 showed an association with a significant expression change (Fig. 5a) (Table S9) (Methods). Most of the significant changes were associated with INTRON SMRs in skin (SKCM) and colorectal (CRC) tumors (Figures S16 and S17). The motifs PTBP1, PCBP1, RBM8A and ZNF638 harbored the highest number of mutations associated with significant transcript expression changes (Fig. 5b) (Figure S17). In particular, mutations in RBM8A motifs were associated with expression changes of the histone acetyl-transferase gene *KAT6A* and the pre-B-cell leukemia transcription factor 1 gene *PBX1*, both potential oncogenes (43,44). In the case of PBX1, two transcript isoforms change expression in opposite directions, indicating an isoform switch.

**Figure 5.**
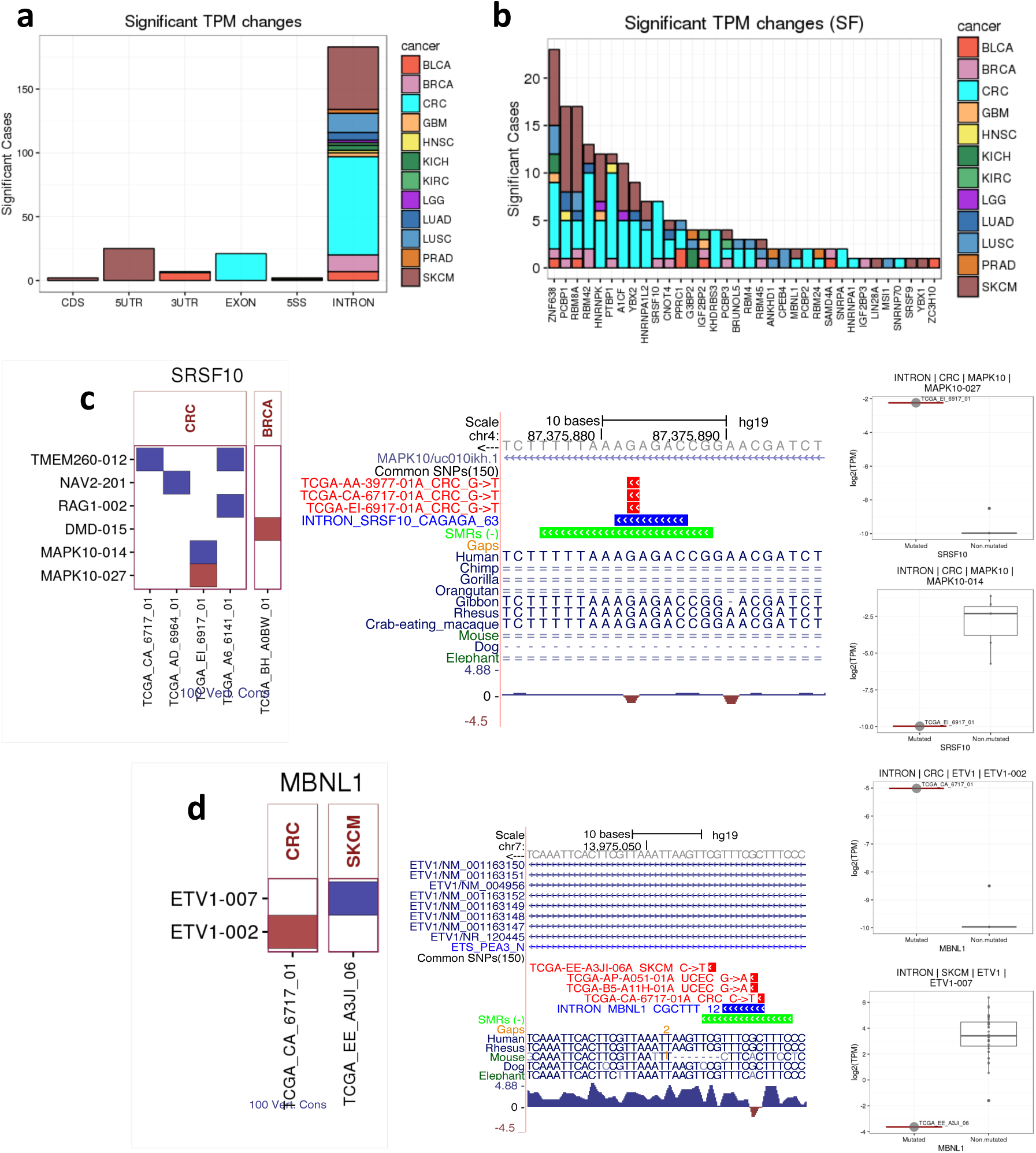
Transcript expression changes associated to mutations in RBP motifs. **(a)** For each region type (x axis) we give the number of SMRs (y axis) for which we found a significant change in transcript isoform expression associated with somatic mutations in enriched motifs within the SMR. The counts are color-coded by tumor type. **(b)** For each motif (x axis), we give the number of cases for which we found a significant change in transcript expression (y axis) associated with somatic mutations in the motif. The counts are color-coded by tumor type. **(c)** Significant changes in transcript expression associated with mutations in intronic SRSF10 motifs. For each patient (x axis) we show the transcripts (y axis) that have a significant increase (red) or decrease (blue) in expression. We separate them according to tumor type (indicated above). In the UCSC screen shot we indicate the patients and mutations on the CAGAGA motif in the intron of *MAPK10*. Below we show the significant expression changes detected in two different transcripts of *MAPK10* associated to mutations in the SRSF10 motif. **(d)** Significant changes in transcript expression associated with mutations in intronic MBNL1 motifs. For each patient (x axis) we show the transcripts (y axis) that have a significant increase (red) or decrease (blue) in expression. We separate them according to tumor type (indicated above). In the UCSC screen shot we indicate the patients and mutations on the CGCTTT motif in the intron of *ETV1*. Below we show the significant expression changes detected in two different transcripts of *ETV1* associated to mutations in the MBNL1 motif.

We also found mutations in an intronic SRSF10 motif associated with expression changes in a transcript from the dystrophin gene (*DMD*) in breast cancer (BRCA), and in two transcripts from the Mitogen-Activated Protein Kinase 10 *MAPK10 (JNK3*) in colorectal cancer (CRC) (Fig. 5c). In *DMD*, the mutation was associated with increased expression, whereas in *MAPK10* we observed an isoform switch (Fig. 5c). *MAPK10* is a pro-apoptotic gene and our results suggest that a mutation in an intronic SRSF10 motif conserved in primates to be associated to an isoform switch in CRC (Fig. 5c). We also found a CRC mutation on an intronic MBNL1 motif conserved in mammals that is associated with an upregulation of the transcription factor ETV1 (Fig. 5d), which has been linked to prostate cancer (45). A mutation at a nearby site in SKCM shows association to downregulation of a different transcript isoform (Fig. 5d). In 5UTR SMRs, significant changes were associated only with SKCM mutations (Figure S18). One of them corresponds to a mutation on a PCBP3 motif associated with an isoform switch in the galactokinase-2 gene *GALK2* in melanoma. *GALK2* is a regulator of prostate cancer cell growth (46) and has alternative splicing in multiple tumor types (29). Our results indicate that these splicing alterations could stem from a mutation on a PCBP3 motif at the 5’UTR. Finally, among the changes associated with 3UTR SMRs, we found a mutation in a HuR motif in CRC related to the downregulation of the Debrin-like gene *DBNL* (Figure S18).

We also analyzed the changes in all possible exon-exon junctions defined from spliced reads mapped to the genome and overlapping SMRs. Of the SMRs tested, 30 were associated with a significant inclusion change in at least one junction (Fig. 6a) (Table S10). The majority of cases occurred in INTRON SMRs, and associated to mutations in bladder (BLCA) and head and neck squamous-cell (HNSC) tumors. Significant junctions were also commonly associated with 5SS SMRs, mainly in uterine (UCEC) and skin (SKCM) tumors (Fig. 6a) and in 5UTRs SMRs specifically associated to SKCM mutations. In contrast, we found few associations for 3SS and EXON SMRs, and none for CDS SMRs (Fig. 6a). Significant changes in junctions were most commonly associated to the 5SSC and 3SSC motifs, as well as to IGF2BP2, HNRNPA2B1, HNRNPLL and PCBP1 motifs among others (Fig. 6b). Among the significant changes associated to 5SSC (Fig. 6c), a mutation in the peptidyl-tRNA hydrolase 2 gene *PTRH2* in uterine cancer (UCEC) would lead to the recognition of an upstream cryptic 5’ss (Fig. 6d). *PTRH2* induces anoikis and its downregulation is linked to metastasis in tumor cells (47). The mutation observed could therefore play a role in cancer progression. We also found a significant splicing change in the Farnesyl Pyrophosphate Synthetase gene *FDPS* in melanoma that would induce an alternative 5’ss and skip part of the Polyprenyl synthetase domain (Figure S19). *FDPS* induces autophagy in cancer cells (48), and an alteration of the autophagy pathway has been related to myeloid neoplasms when *SF3B1* mutations are present (49). It would be interesting to investigate further whether this splicing-induced alteration of *FDPS* could recapitulate a similar phenotype. Among the significant changes associated with 3SS mutations there was one in the Adenine Phosphoribosyl transferase gene *APRT* associated to G>A mutations at the consensus 3’ss splice site in melanoma (SKCM), which would skip the Phosphoribosyl transferase domain (Figure S19).

**Figure 6.**
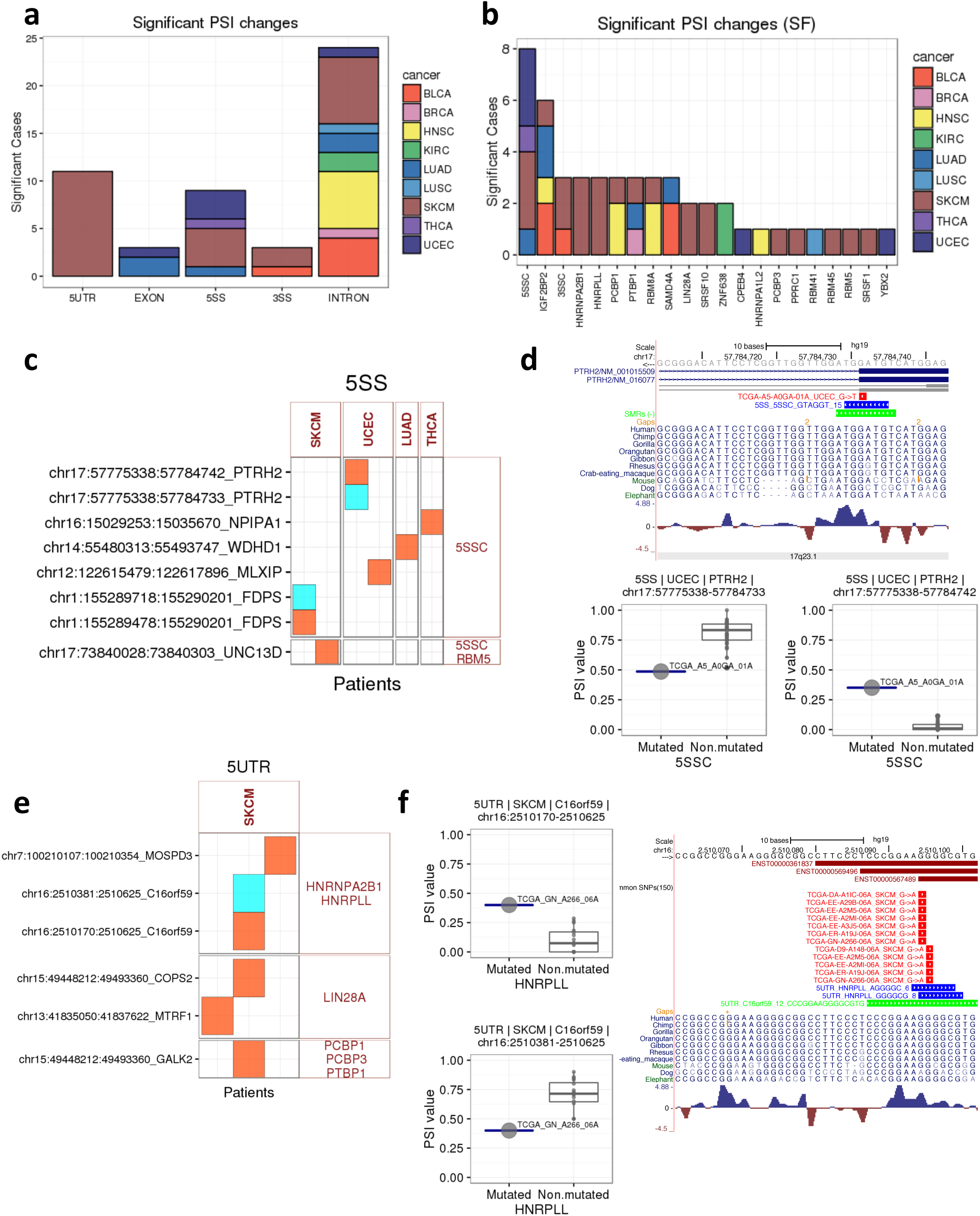
Changes in junction usage associated to mutations in RBP motifs. **(a)** For each region type (x axis) we give the number of SMRs (y axis) for which we found a significant change in exon-exon junction inclusion associated with somatic mutations in enriched motifs within the SMR. The counts are color-coded by tumor type. **(b)** For each motif (x axis), we give the number of instances (y axis) for which we found a significant change in exon-exon junction inclusion associated with somatic mutations in the motif (x axis). The counts are color-coded by tumor type. **(c)** Significant changes in junction inclusion associated to mutations in 5’ splice site (5’ss) motifs. For each patient (x axis) we show the junctions (y axis) that have a significant increase (orange) or decrease (cyan) in inclusion (PSI). We separate them according to tumor type (indicated above). **(d)** Significant junction changes in *PTRH2* in uterine cancer (UCEC). We show the changing junctions in gray. The mutation in the annotated 5’ss induces the usage of an upstream cryptic 5’ss. We show the SMR (green), the enriched motif (blue), and the mutation (red). The boxplots blow show the PSI values (y axis) of the two changing junctions separated by samples with mutations and without mutations in this SMR in UCEC, indicated in the x axis. **(e)** Significant changes in junction inclusion associated to mutations in 5UTR SMRs. For each patient (x axis) we show the junctions (y axis) that have a significant increase (orange) or decrease (cyan) in inclusion (PSI). We separate them according to tumor type (indicated above). **(f)** Significant junction changes in *C16orf59* associated to mutations in an HRNPLL motif in melanoma (SKCM). The boxplots show the PSI values (y axis) of the two changing junctions separated by samples with mutations and without mutations, indicated in the x axis. In the screenshot we show the 5UTR SMR (green), the enriched motifs (blue), and the mutations (red), which suggest dinucleotide mutations GG>AA in some patients. The junctions associated to these mutations are downstream of the SMR and do not appear in the genomic range shown in the figure.

We also found mutations in 5UTR and EXON motifs associated to splicing changes (Fig. 6e). We found a mutation in a 5UTR HNRPLL motif in *C16orf59* that is conserved across mammals (Fig. 6f). We also found a mutation in an EXON IGF2BP2 motif in the *C14orf37* locus (Figure S19). Finally, INTRON SMRs had the most numerous number of junction changes associated with mutations in multiple RBP motifs, including RBM8A and SRSF10 (Figure S20), which included changes in the histone gene *HIST1H2AC* related to mutations in the RBM8A motif in HNSC, and in *GMEB1* related to mutations in an SRSF10 motif. BED and GFF tracks representing all the found cases are available as supplementary material.

## Discussion

We have described a novel method to identify and characterize somatic mutations in coding and non-coding regions in relation to their potential to be involved in protein-RNA binding. By considering mutational significance in combination with the enrichment of RBP binding motifs, we identified mutations in non-coding regions, particularly in deep intronic regions, with evidence of impact on RNA processing. Our analysis provides new potential mechanisms by which somatic mutations impact RNA processing and contribute to tumor phenotypes. Although these mutations may not be as frequent as other classical oncogenic mutations, recent evidence suggests that rare somatic variants can have an impact on expression and be clinically actionable (50). As RBPs are known to control entire cellular pathways, from epithelial-to-mesenchymal transition (51) to cellular differentiation (52), these results suggest a general model by which cancer mutations disrupt the function of RBP-targets and contribute independently to the disruption of similar pathways (Fig. 7). Our work also provides a new strategy to interpret non-coding mutations and indicates that lowly recurrent mutations could still be relevant to the study of cancer, as they can impact functions collectively controlled by the same RBP.

**Figure 7.**
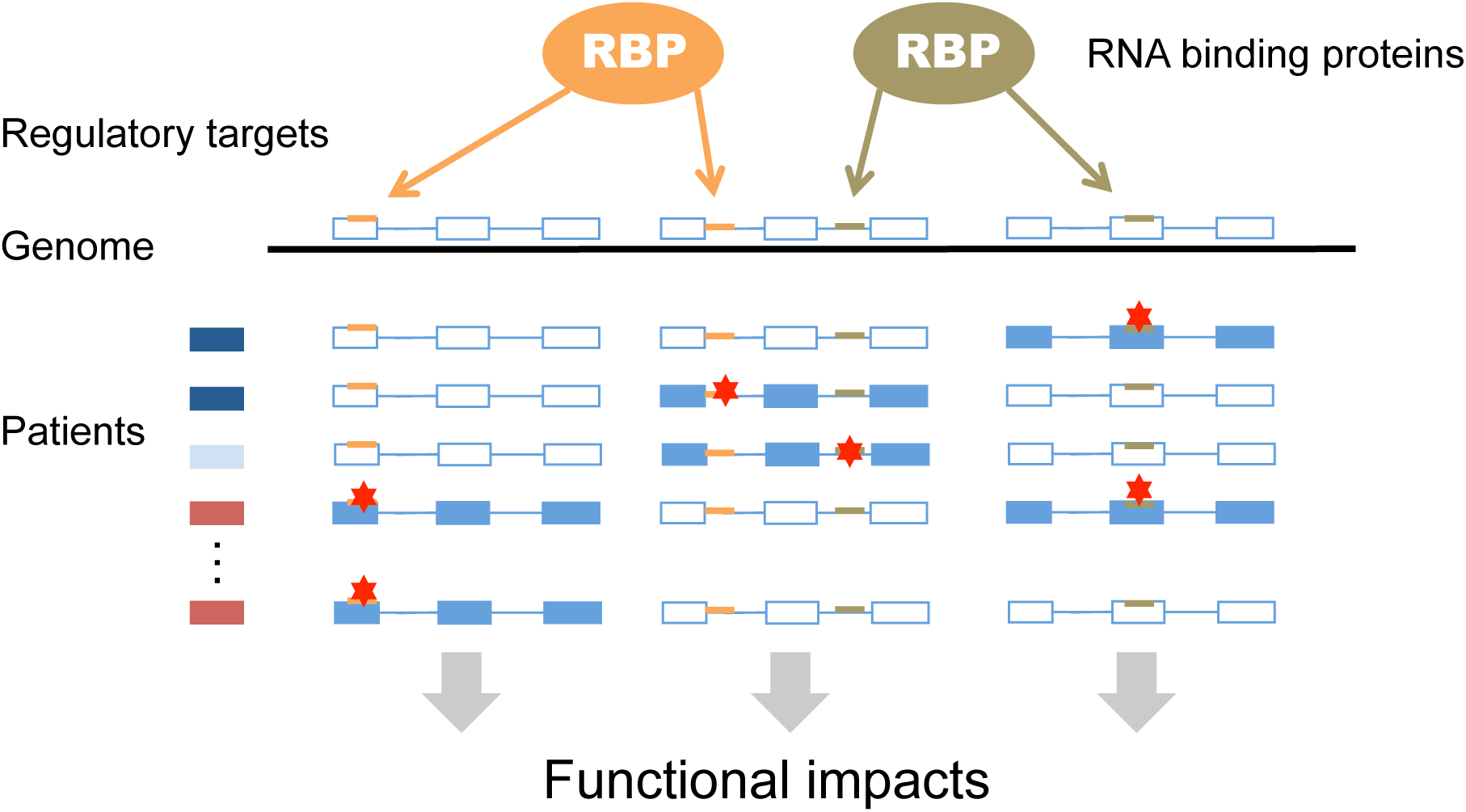
Non-coding mutations in human tumors impact binding sites of RNA binding proteins. Our analysis suggests that many of the mutations (indicated in red) on non-coding regions, predominantly introns, and UTRs, impact binding sites of RNA binding proteins (indicated in orange and green) and affect RNA processing in multiple different genes across patients. In the figure altered RNA processing are indicated as solid gene models. These alterations would contribute to the frequent changes observed in RNA processing in tumors and could indicate novel oncogenic mechanisms.

MIRA presents several advantages with respect to previous methods. It detects deep intronic mutations, whereas previous methods only tested intronic regions immediately adjacent to exons (11,15). Additionally, unlike previous approaches (11,13,15), we examined the location of the mutations in the context of a RNA regulatory motif, which enables the combined interpretation of non-coding mutations at multiple sites. Additionally, our analyses were driven by the positions of mutations on regulatory motifs, hence providing the precise region where mutations likely play a role. Finally, unlike most previous approaches, we tested the impact on RNA processing and expression using RNA-seq from the same samples. We observed significant changes for a small fraction of SMRs, indicating that the overall impact on RNA is modest as measured by RNA sequencing from cancer tissue samples. Deeper sequencing of tissue samples or in-vitro based assays could help validating many more of these events.

The majority of the validated cases were deep-intronic mutations, whose interpretation has remained elusive so far. Here we provided evidence of their potential relevance in cancer. Interestingly, whereas other methods assumed that the function of lncRNAs may be impacted only through exonic mutations (15), we found that intronic SMRs may be relevant as well. Additionally, although it has been generally assumed that RNA structure determines the function in UTR regions (11), we found 5UTR SMRs associated to RNA processing changes, indicating new mechanisms. Our approach is subject to several limitations. The analysis may be underpowered due to the relatively small number of patients analyzed. Another limitation is that we chose specific descriptions for the RNA binding motifs; hence the analysis is limited by their accuracy. Despite computational and technological advances, precise definitions of RBP binding sites at a genome scale remains challenging, and different RBPs may bind similar sequence motifs. To deal with these ambiguities, our method allowed for multiple assignments of 6-mers to RBP labels, providing the opportunity to describe the mutational patterns of different RBP motifs. In summary, MIRA analyzes non-coding mutations to take into account functional analogous sites at different genomic positions and RNA-related selection processes, providing evidence that multiple RNA processing mechanisms may be impaired in cancer through mutations on RBP motifs, thereby uncovering novel alterations relevant for cancer.

## Methods

### Data

We used the Gencode gene annotations (v19), excluding pseudogenes. To define gene loci unequivocally, we clustered transcripts that shared a splice site on the same strand, and considered a gene to be the genomic locus and the strand defined by those transcripts. We used somatic mutations from whole genome sequencing for 505 tumor samples from 14 tumor types published previously (Table S1) (10): bladder carcinoma (BLCA) (21 samples), breast carcinoma (BRCA) (96 samples), colorectal carcinoma (CRC) (42 samples), glioblastoma multiforme (GBM) (27 samples), head and neck squamous carcinoma (HNSC) (27 samples), kidney chromophobe (KICH) (15 samples), kidney renal carcinoma (KIRC) (29 samples), low grade glioma (LGG) (18 samples), lung adenocarcinoma (LUAD) (46 samples), lung squamous cell carcinoma (LUSC) (45 samples), prostate adenocarcinoma (PRAD) (20 samples), skin carcinoma (SKCM) (38 samples), thyroid carcinoma (THCA) (34 samples), and uterine corpus endometrial carcinoma (UCEC) (47 samples). We only used substitutions, discarding those with a precise allelic match to a germline variant in dbSNP138.

### Detection of significantly mutated regions

To identify significantly mutated regions (SMRs) in both coding and non-coding regions of genes we used a sliding-window approach, whereby along each gene locus we tested all overlapping windows of length 7 (7-mer window) that harbored at least one mutation. For each 7-mer window, we performed a double statistical test. First, given a window with *n* mutations in a gene of length *L* and *N* mutations overall, we performed a binomial test using *N/L* as the expected local mutation rate (Figure S1). All tested windows in a gene were adjusted for multiple testing using the Benjamini-Hochberg (BH) method, and we kept only windows with false discovery rate (corrected p-value) < 0.05 (Figure S2).

To account for potential nucleotide biases we performed a second test per 7-mer window: we compared the mutation count in a given window with the expected count according to the distribution of mutations per nucleotide in the same gene as follows: For each base *a* we calculated the rate of mutations in a locus *R(a) = m(a)/n(a*), where *n(a*) is the number of *a* bases in the gene and *m(a*) is the number of those bases that are mutated. The expected mutation count is then calculated using the nucleotide counts in the window and the mutation rate per nucleotide. For instance, for the 7-mer window AACTGCAG, the expected count was calculated as: *E = 3R(A) + 2R(C) + 2R(G) + R(T*). This was compared to the actual number of mutations, *n*, observed in that window to define a nucleotide bias (NB) score: *NB-score* = *log2(n/E*) per window. We discarded windows corresponding to single-nucleotide repeats (e.g. AAAAAAA) and kept windows with NB-score > 6 (Figure S2). Further, we kept 7-mer windows that overlapped any of the three intronic or exonic bases around exon-intron boundaries with 1 or more mutations as long as the NB-score was greater than 6.

Significant 7-mer windows were clustered according to genomic overlap and classified according to the genic region in the same strand on which they fell: 5’ or 3’ untranslated regions (5UTR/3UTR), coding sequence (CDS), exon in short or long non-coding RNA (EXON), 5’/3’ splice-site (5SS/3SS), or intron (INTRON). To unambiguously assign each 7-mer window we prioritized regions as follows: 5SS/3SS > CDS > 5UTR/3UTR > EXON > INTRON, i.e. if a window overlapped a splice-site, it was classified as such; else, if it overlapped a CDS, it was classified as CDS; etc. No significant window overlapped start or stop codons. To each SMR we assigned the average NB-score and a corrected p-value using the Simes approach (53): we ranked the p-values of the n overlapping windows in increasing order *p*_*i*_, *i=1,2,…n* and calculated *p*_*s*_ = *min*{ *np_1_ / 1, np_2_ / 2, np_3_ / 3, …., np*_*n*_ / *n*}, where *p*_1_ was the lowest and *p_n_* the highest p-values in the cluster. Each SMR cluster was then assigned the p-value *p_s_*. Using other window lengths (k=6,8,9) we observed that k=7 provides an optimal trading off between the number of regions covered and the clustering of those regions (Figure S2). Code for this analysis is available at https://github.com/comprna/mira.

### Comparison to expression, replication timing and to other methods

Data for replication time was obtained from (14). Only SMRs with replication time data were analyzed. For each SMR in the PAN505 cohort, we considered annotated transcripts whose genomic sequence overlapped with an SMR. We calculated the total expression in transcripts per million (TPM) units for the overlapping transcripts per patient and averaged them across patients. For each SMR we compared the average expression of the SMR-containing transcripts in the mutated samples with the number of mutations. Additionally, using the same mutation dataset, we analyzed all SMRs with LARVA (14). LARVA assesses the significance of mutations in any genomic region integrating multiple non-coding functional elements, modeling their mutation count with a beta-binomial distribution to handle over dispersion related to the mutation heterogeneity and mutation correlation between neighboring sites, and uses regional genomic features such as replication timing to better estimate local mutation rates and mutational enrichments. We compared the significance of our SMRs with the significance given by LARVA using the model with a beta-binomial distribution and the replication timing correction (p-bbd-cor), which accounts for over dispersion of the mutation rates and regional biases. We also analyzed our SMRs with OncodriveFML (11), which evaluates the functional impact of mutations using CADD score (54) and RNA secondary structure (55), using the same mutation dataset. OncodriveFML was run with option *—type coding* for SMRs of type CDS, and with the option *—type noncoding* for all other SMR types (5SS/3SS, 5UTR, 3UTR, INTRON, EXON, where EXON corresponds to exons from non-coding RNAs). Each type was run independently.

### Control regions for SMR comparison

For each SMR, we sampled 100 random regions of the same length and same type from the Gencode annotation, without mutations, and allowing for a maximum variation of G+C content of 5%. Each of these 100 controls was separated into different sets to generate 100 control sets of the same number as SMRs, each with similar distribution and G+C content distributions. For the 5SS and 3SS SMRs we generated controls by sampling regions with the same length and same relative position from Gencode exon-intron boundaries without mutations and controlling for G+C content.

### Motif analysis

We performed an unbiased search for enriched k-mers (k=6) on the SMRs using MoSEA (https://github.com/comprna/MoSEA) (29), using as input the SMR sequences and control regions, for each region type. MoSEA counts the number of SMRs and control regions in which each 6-mer appears and a z-score was computed for each 6-mer comparing the observed frequency with the distribution of frequencies in 100 control subsamples of the same size, length distribution and GC content as the SMR set. This was repeated reversing the strand of SMRs and control regions, and those 6-mers that appeared significantly enriched in the direct and reversed analyses for the same region type were discarded. We considered significant the 6-mers with z-score > 1.96 and 5 or more counts. This analysis included the GT and AG containing 6-mers at 5SS and 3SS SMRs, respectively. In total we obtained 357 enriched 6-mers (Table S4) in 3547 SMRs (Table S5). From all 20307 SMRs, 749 (3,68%) appeared in both strands due to overlapping genes. However, considering the 3456 SMRs with enriched 6-mers, only 25 (0,72%) overlapped on opposite strands (Fisher’s exact test p-value = 1.29e-32, odds-ratio = 6.16). To label enriched 6-mers we used DeepBind (30) to score each 6-mer using models for 522 proteins containing KH (24 proteins), RRM (134, 2 in common with KH) and C2H2 (366 proteins) domains from human (413 proteins), mouse (49 proteins) and Drosophila (60 proteins). For each 6-mer, we kept the top three predictions with score > 0.1. Subsequently, given all 6-mers associated to the same RBP label, we kept those 6-mers at a maximum Levenshtein distance of 2 from the top-scoring 6-mer. Levenshtein distance measures the dissimilarity between two strings based on the number of deletions, insertions or substitutions required to transform one string into the other.

### Significant mutations per position of a motif

For each RBP we considered all the associated 6-mers in SMRs of a given region type and performed a multiple sequence alignment (MSA) using ClustalW (56). Sequence logos were built from this alignment and somatic mutations were counted per position relative to the MSA. As a control, we shuffled the mutations along the aligned positions to calculate an average per position. Germline mutations per position of the MSA from the 1000 genomes project (57) were also considered.

### Gene set enrichment analysis

Annotations for gene sets were obtained from the Molecular Signatures Database v4.0 (58). We performed a Fisher’s exact test per hallmark set for genes harboring SMRs with labeled enriched motifs using the counts of genes with/without the RBP motif SMRs, and within/outside each gene set.

### CLIP-Seq analysis

We collected binding sites from 91 CLIP-Seq experiments for 68 different RBPs from multiple sources (34–40) and selected the available significant CLIP clusters from each experiment. For datasets with two replicates we selected the intersecting regions of the significant CLIP clusters, i.e. the genomic ranges covered by both replicates. From all SMRs with assigned RBP label, 482 (13.6%) had CLIP signal, whereas 3064 did not have CLIP signal. In contrast, from all SMRs without assigned RBP label, 1239 had CLIP overlap (7.4%), whereas 15522 did not have CLIP signal (Fisher’s exact test p-value p-value = 4.10e-30, odds-ratio: 1.97). We performed the same analysis per RBP label: For each RBP label, we considered the SMRs with and without the label assignment and tested the enrichment of CLIP signal associated to the label using a Fisher’s exact test (Table S7) (Figure S10).

### RNA-seq data analysis

TCGA RNA-seq data was obtained for the PAN505 samples from the Genomic Data Commons (https://gdc-portal.nci.nih.gov/). We estimated transcript abundances for the Gencode annotations (v19) in TPM units using Salmon (Version 0.8.1) (59). For each mutated position in an enriched motif, we calculated the association to a transcript expression change using an outlier statistic. For each transcript whose genomic extension contained the SMR with the mutated motif, we compared the transcript log_2_(TPM+0.01) for each patient with the mutation, with the distribution of log_2_(TPM+0.01) values for the same transcript in the patients from the same tumor type with no mutations in the SMR. We only considered those cases where at least 5 patients lacked a mutation. We kept those cases with |z-score|>1.96 and a difference between the observed log_2_(TPM+0.01) and the mean of log_2_(TPM+0.1) in patients without mutations greater than 0.5 in absolute value, and considered significant those with a p-value < 0.05 after adjusting for multiple testing (BH approach). RNA-seq reads were also mapped to the human genome (hg19) with STAR (version 2.5.0) (60) and analyzed using Junckey (https://github.com/comprna/Junckey) as follows. All exon-exon junctions defined by spliced reads that appeared in any of the samples were grouped into junction-clusters. Any two junctions were placed in the same cluster if they shared at least one splice-site. Clusters were built using all junctions present in any patient, but junction read-counts were assigned per patient. Only clusters with at least 30 reads in all samples were used. Additionally, we only used junctions <100kbp in length and with >1% of reads from the cluster in all samples. For each patient, the read-count per junction was normalized by the total read count in that cluster to define the junction inclusion or proportion spliced-in (PSI). A junction not expressed in a sample was set to PSI=0. For each SMR containing an enriched motif, we compared each patient with a mutation in the motif against all patients for the same tumor type without mutations in the same SMR. We measured a z-score derived from the PSI for each junction overlapping the motif and the of PSIs for the same junction in the non-mutated patients, and we kept only those cases with |z-score|>1.96 and |ΔPSI|>0.1. We considered significant those changes with p-value < 0.05 after adjusting for multiple testing (BH approach).

## Supplementary Data and Software

UCSC tracks for SMRs, mutations, motifs and differentially included junctions are given as supplementary file. Plots for all mutated RBP binding motifs and RNA processing changes are available at: http://comprna.upf.edu/Data/MutationsRBPMotifs/ Code used in this manuscript is available at:

https://github.com/comprna/mira

https://github.com/comprna/MoSEA

https://github.com/comprna/Junckey

## Acknowledgements

We thank M. Taylor, E. Larsson and N. Lopez-Bigas for discussions. The results published here are in whole or part based upon data generated by The Cancer Genome Atlas managed by the NCI and NHGRI. Information about TCGA can be found at http://cancergenome.nih.gov.

## List of Abbreviations

SMR: significantly mutated region
NB-score: Nucleotide-bias score
RBP: RNA binding protein
RNA-seq: RNA sequencing;

## Authors’ contributions

EE proposed and led the study. BS implemented the methods and performed the analyses. JLT developed the method for junction analysis. PJT and SRP performed the mapping of RNA-seq reads and transcript quantification. EE and BS wrote the manuscript with essential inputs from JLT, PJT and SRP.

## References

1. Hanahan D, Weinberg RA. Hallmarks of Cancer: The Next Generation. Cell. 2011;144:646–74.

2. Vogelstein B, Papadopoulos N, Velculescu VE, Zhou S, Diaz Jr. LA, Kinzler KW. Cancer Genome Landscapes. Science (80-). 2013;339:1546–58.

3. Alexandrov LB, Nik-Zainal S, Wedge DC, Campbell PJ, Stratton MR. Deciphering signatures of mutational processes operative in human cancer. Cell Rep. 2013;3:246–59.

4. Weinhold N, Jacobsen A, Schultz N, Sander C, Lee W. Genome-wide analysis of noncoding regulatory mutations in cancer. Nat Genet [Internet]. Nature Publishing Group; 2014;46:1160–5. Available from: http://www.pubmedcentral.nih.gov/articlerender.fcgi?artid=4217527&tool=pmcentrez&rendertype=abstract

5. Juul M, Bertl J, Guo Q, Nielsen MM, Świtnicki M, Hornshøj H, et al. Non-coding cancer driver candidates identified with a sample-and position-specific model of the somatic mutation rate. Elife [Internet]. 2017;6. Available from: http://www.ncbi.nlm.nih.gov/pubmed/28362259

6. Horn S, Figl A, Rachakonda PS, Fischer C, Sucker A, Gast A, et al. TERT Promoter Mutations in Familial and Sporadic Melanoma. Science (80-). 2013;339:959–61.

7. Huang FW, Hodis E, Xu MJ, Kryukov G V., Chin L, Garraway LA. Highly Recurrent TERT Promoter Mutations in Human Melanoma. Science (80-) [Internet]. 2013 [cited 2017 Jan 28];339:957–9. Available from: http://www.ncbi.nlm.nih.gov/pubmed/23348506

8. Piraino SW, Furney SJ. Beyond the exome: the role of non-coding somatic mutations in cancer. Ann Oncol Off J Eur Soc Med Oncol [Internet]. 2016;27:240–8. Available from: http://www.ncbi.nlm.nih.gov/pubmed/26598542

9. Melton C, Reuter JA, Spacek D V, Snyder M. Recurrent somatic mutations in regulatory regions of human cancer genomes. Nat Genet [Internet]. 2015;47:710–6. Available from: http://www.ncbi.nlm.nih.gov/pubmed/26053494

10. Fredriksson NJ, Ny L, Nilsson JA, Larsson E. Systematic analysis of noncoding somatic mutations and gene expression alterations across 14 tumor types. Nat Genet [Internet]. 2014;46:1–7. Available from: http://dx.doi.org/10.1038/ng.3141

11. Mularoni L, Sabarinathan R, Deu-Pons J, Gonzalez-Perez A, López-Bigas N. OncodriveFML: a general framework to identify coding and non-coding regions with cancer driver mutations. Genome Biol [Internet]. 2016;17:128. Available from: http://genomebiology.biomedcentral.com/articles/10.1186/s13059-016-0994-0

12. Khurana E, Fu Y, Chakravarty D, Demichelis F, Rubin MA, Gerstein M. Role of non-coding sequence variants in cancer. Nat Rev Genet. 2016;17:93–108.

13. Piraino SW, Furney SJ. Identification of coding and non-coding mutational hotspots in cancer genomes. BMC Genomics [Internet]. 2017;18:17. Available from: http://www.ncbi.nlm.nih.gov/pubmed/28056774

14. Lochovsky L, Zhang J, Fu Y, Khurana E, Gerstein M. LARVA: an integrative framework for large-scale analysis of recurrent variants in noncoding annotations. Nucleic Acids Res [Internet]. 2015;43:8123–34. Available from: http://www.ncbi.nlm.nih.gov/pubmed/26304545

15. Lanzós A, Carlevaro-Fita J, Mularoni L, Reverter F, Palumbo E, Guigó R, et al. Discovery of Cancer Driver Long Noncoding RNAs across 1112 Tumour Genomes: New Candidates and Distinguishing Features. Sci Rep [Internet]. 2017;7:41544. Available from: http://www.ncbi.nlm.nih.gov/pubmed/28128360

16. Rissland OS. The organization and regulation of mRNA-protein complexes. Wiley Interdiscip Rev RNA [Internet]. 2017;8. Available from: http://www.ncbi.nlm.nih.gov/pubmed/27324829

17. Ule J, Jensen KB, Ruggiu M, Mele A, Ule A, Darnell RB. CLIP identifies Nova-regulated RNA networks in the brain. Science [Internet]. 2003;302:1212–5. Available from: http://www.ncbi.nlm.nih.gov/pubmed/14615540

18. Lambert N, Robertson A, Jangi M, McGeary S, Sharp PA, Burge CB. RNA Bind-n-Seq: quantitative assessment of the sequence and structural binding specificity of RNA binding proteins. Mol Cell [Internet]. 2014;54:887–900. Available from: http://www.ncbi.nlm.nih.gov/pubmed/24837674

19. Ray D, Kazan H, Cook KB, Weirauch MT, Najafabadi HS, Li X, et al. A compendium of RNA-binding motifs for decoding gene regulation. Nature [Internet]. 2013;499:172–7. Available from: http://www.ncbi.nlm.nih.gov/pubmed/23846655

20. Haerty W, Ponting CP. Unexpected selection to retain high GC content and splicing enhancers within exons of multiexonic lncRNA loci. RNA [Internet]. 2015;21:333–46. Available from: http://www.ncbi.nlm.nih.gov/pubmed/25589248

21. Paronetto MP, Bernardis I, Volpe E, Bechara E, Sebestyén E, Eyras E, et al. Regulation of FAS exon definition and apoptosis by the ewing sarcoma protein. Cell Rep. 2014;7.

22. Soemedi R, Cygan KJ, Rhine CL, Wang J, Bulacan C, Yang J, et al. Pathogenic variants that alter protein code often disrupt splicing. Nat Genet [Internet]. 2017; Available from: http://www.ncbi.nlm.nih.gov/pubmed/28416821

23. Jung H, Lee D, Lee J, Park D, Kim YJ, Park W-Y, et al. Intron retention is a widespread mechanism of tumor-suppressor inactivation. Nat Genet [Internet]. 2015;47:1242–8. Available from: http://dx.doi.org/10.1038/ng.3414

24. Supek F, Miñana B, Valcárcel J, Gabaldón T, Lehner B. Synonymous mutations frequently act as driver mutations in human cancers. Cell [Internet]. 2014;156:1324–35. Available from: http://www.ncbi.nlm.nih.gov/pubmed/24630730

25. Ke S, Shang S, Kalachikov SM, Morozova I, Yu L, Russo JJ, et al. Quantitative evaluation of all hexamers as exonic splicing elements. Genome Res [Internet]. 2011;21:1360–74. Available from: http://www.ncbi.nlm.nih.gov/pubmed/21659425

26. Julien P, Miñana B, Baeza-Centurion P, Valcárcel J, Lehner B. The complete local genotype-phenotype landscape for the alternative splicing of a human exon. Nat Commun [Internet]. 2016;7:11558. Available from: http://www.ncbi.nlm.nih.gov/pubmed/27161764

27. Lawrence MS, Stojanov P, Polak P, Kryukov G V, Cibulskis K, Sivachenko A, et al. Mutational heterogeneity in cancer and the search for new cancer-associated genes. Nature [Internet]. 2013;499:214–8. Available from: http://www.ncbi.nlm.nih.gov/pubmed/23770567

28. Liu L, De S, Michor F. DNA replication timing and higher-order nuclear organization determine single-nucleotide substitution patterns in cancer genomes. Nat Commun [Internet]. 2013;4:1502. Available from: http://www.ncbi.nlm.nih.gov/pubmed/23422670

29. Sebestyén E, Singh B, Miñana B, Pagès A, Mateo F, Pujana MA, et al. Large-scale analysis of genome and transcriptome alterations in multiple tumors unveils novel cancer-relevant splicing networks. Genome Res [Internet]. 2016;26:732–44. Available from: http://www.ncbi.nlm.nih.gov/pubmed/27197215

30. Alipanahi B, Delong A, Weirauch MT, Frey BJ. Predicting the sequence specificities of DNA-and RNA-binding proteins by deep learning. Nat Biotechnol [Internet]. 2015;33:831–8. Available from: www.ncbi.nlm.nih.gov/pubmed/26213851

31. Chabot B, Shkreta L. Defective control of pre-messenger RNA splicing in human disease. J Cell Biol [Internet]. 2016;212:13–27. Available from: http://www.ncbi.nlm.nih.gov/pubmed/26728853

32. Tripathi V, Sixt KM, Gao S, Xu X, Huang J, Weigert R, et al. Direct Regulation of Alternative Splicing by SMAD3 through PCBP1 Is Essential to the Tumor-Promoting Role of TGF-?. Mol Cell [Internet]. 2016;64:549–64. Available from: http://www.ncbi.nlm.nih.gov/pubmed/27746021

33. Oberdoerffer S, Moita LF, Neems D, Freitas RP, Hacohen N, Rao A. Regulation of CD45 alternative splicing by heterogeneous ribonucleoprotein, hnRNPLL. Science [Internet]. 2008;321:686–91. Available from: http://www.ncbi.nlm.nih.gov/pubmed/18669861

34. Sundararaman B, Zhan L, Blue SM, Stanton R, Elkins K, Olson S, et al. Resources for the Comprehensive Discovery of Functional RNA Elements. Mol Cell [Internet]. 2016;61:903–13. Available from: http://www.ncbi.nlm.nih.gov/pubmed/26990993

35. Bechara EG, Sebestyén E, Bernardis I, Eyras E, Valcárcel J. RBM5, 6, and 10 differentially regulate NUMB alternative splicing to control cancer cell proliferation. Mol Cell [Internet]. 2013;52:720–33. Available from: http://www.ncbi.nlm.nih.gov/pubmed/24332178

36. Raj B, Irimia M, Braunschweig U, Sterne-Weiler T, O’Hanlon D, Lin ZY, et al. A global regulatory mechanism for activating an exon network required for neurogenesis. Mol Cell. 2014;56:90–103.

37. Shao C, Yang B, Wu T, Huang J, Tang P, Zhou Y, et al. Mechanisms for U2AF to define 3’ splice sites and regulate alternative splicing in the human genome. Nat Struct Mol Biol [Internet]. 2014;21:997–1005. Available from: http://www.ncbi.nlm.nih.gov/pubmed/25326705

38. Rodor J, Pan Q, Blencowe BJ, Eyras E, Cáceres JF. The RNA-binding profile of Acinus, a peripheral component of the exon junction complex, reveals its role in splicing regulation. RNA [Internet]. 2016;22:1411–26. Available from: http://www.ncbi.nlm.nih.gov/pubmed/27365209

39. Best A, James K, Dalgliesh C, Hong E, Kheirolahi-Kouhestani M, Curk T, et al. Human Tra2 proteins jointly control a CHEK1 splicing switch among alternative and constitutive target exons. Nat Commun [Internet]. 2014;5:4760. Available from: http://www.ncbi.nlm.nih.gov/pubmed/25208576

40. Yang Y-CT, Di C, Hu B, Zhou M, Liu Y, Song N, et al. CLIPdb: a CLIP-seq database for protein-RNA interactions. BMC Genomics [Internet]. 2015;16:51. Available from: http://www.ncbi.nlm.nih.gov/pubmed/25652745

41. Sawicka K, Bushell M, Spriggs KA, Willis AE. Polypyrimidine-tract-binding protein: a multifunctional RNA-binding protein. Biochem Soc Trans [Internet]. 2008;36:641–7. Available from: http://www.ncbi.nlm.nih.gov/pubmed/18631133

42. Pichon X, Wilson LA, Stoneley M, Bastide A, King HA, Somers J, et al. RNA binding protein/RNA element interactions and the control of translation. Curr Protein Pept Sci [Internet]. 2012;13:294–304. Available from: http://www.ncbi.nlm.nih.gov/pubmed/22708490

43. Turner-Ivey B, Guest ST, Irish JC, Kappler CS, Garrett-Mayer E, Wilson RC, et al. KAT6A, a chromatin modifier from the 8p11-p12 amplicon is a candidate oncogene in luminal breast cancer. Neoplasia [Internet]. 2014;16:644–55. Available from: http://www.ncbi.nlm.nih.gov/pubmed/25220592

44. Magnani L, Ballantyne EB, Zhang X, Lupien M. PBX1 genomic pioneer function drives ER? signaling underlying progression in breast cancer. PLoS Genet [Internet]. 2011;7:e1002368. Available from: http://www.ncbi.nlm.nih.gov/pubmed/22125492

45. Cai C, Hsieh C-L, Omwancha J, Zheng Z, Chen S-Y, Baert J-L, et al. ETV1 is a novel androgen receptor-regulated gene that mediates prostate cancer cell invasion. Mol Endocrinol [Internet]. 2007;21:1835–46. Available from: http://www.ncbi.nlm.nih.gov/pubmed/17505060

46. Whitworth H, Bhadel S, Ivey M, Conaway M, Spencer A, Hernan R, et al. Identification of kinases regulating prostate cancer cell growth using an RNAi phenotypic screen. PLoS One [Internet]. 2012;7:e38950. Available from: http://www.ncbi.nlm.nih.gov/pubmed/22761715

47. Karmali PP, Brunquell C, Tram H, Ireland SK, Ruoslahti E, Biliran H. Metastasis of tumor cells is enhanced by downregulation of Bit1. PLoS One [Internet]. 2011;6:e23840. Available from: http://www.ncbi.nlm.nih.gov/pubmed/21886829

48. Wasko BM, Dudakovic A, Hohl RJ. Bisphosphonates induce autophagy by depleting geranylgeranyl diphosphate. J Pharmacol Exp Ther [Internet]. 2011;337:540–6. Available from: http://www.ncbi.nlm.nih.gov/pubmed/21335425

49. Visconte V, Przychodzen B, Han Y, Nawrocki ST, Thota S, Kelly KR, et al. Complete mutational spectrum of the autophagy interactome: a novel class of tumor suppressor genes in myeloid neoplasms. Leukemia [Internet]. 2017;31:505–10. Available from: http://www.ncbi.nlm.nih.gov/pubmed/27773925

50. Berger AH, Brooks AN, Wu X, Shrestha Y, Chouinard C, Piccioni F, et al. High-throughput Phenotyping of Lung Cancer Somatic Mutations. Cancer Cell [Internet]. 2016;30:214–28. Available from: http://www.ncbi.nlm.nih.gov/pubmed/27478040

51. Shapiro IM, Cheng AW, Flytzanis NC, Balsamo M, Condeelis JS, Oktay MH, et al. An emt-driven alternative splicing program occurs in human breast cancer and modulates cellular phenotype. PLoS Genet. 2011;7.

52. Han H, Irimia M, Ross PJ, Sung H-K, Alipanahi B, David L, et al. MBNL proteins repress ES-cell-specific alternative splicing and reprogramming. Nature [Internet]. 2013;498:241–5. Available from: http://www.pubmedcentral.nih.gov/articlerender.fcgi?artid=3933998&tool=pmcentrez&rendertype=abstract

53. Lun ATL, Smyth GK. De novo detection of differentially bound regions for ChIP-seq data using peaks and windows: controlling error rates correctly. Nucleic Acids Res [Internet]. 2014;42:e95. Available from: http://www.ncbi.nlm.nih.gov/pubmed/24852250

54. Kircher M, Witten DM, Jain P, O’Roak BJ, Cooper GM, Shendure J. A general framework for estimating the relative pathogenicity of human genetic variants. Nat Genet [Internet]. 2014;46:310–5. Available from: http://www.ncbi.nlm.nih.gov/pubmed/24487276

55. Sabarinathan R, Tafer H, Seemann SE, Hofacker IL, Stadler PF, Gorodkin J. RNAsnp: efficient detection of local RNA secondary structure changes induced by SNPs. Hum Mutat [Internet]. 2013;34:546–56. Available from: http://www.ncbi.nlm.nih.gov/pubmed/23315997

56. Thompson JD, Gibson TJ, Higgins DG. Multiple sequence alignment using ClustalW and ClustalX. Curr Protoc Bioinforma [Internet]. 2002;Chapter 2:Unit 2.3. Available from: http://www.ncbi.nlm.nih.gov/pubmed/18792934

57. 1000 Genomes Project Consortium, Abecasis GR, Altshuler D, Auton A, Brooks LD, Durbin RM, et al. A map of human genome variation from population-scale sequencing. Nature [Internet]. 2010;467:1061–73. Available from: http://www.ncbi.nlm.nih.gov/pubmed/20981092

58. Liberzon A, Birger C, Thorvaldsdóttir H, Ghandi M, Mesirov JP, Tamayo P. The Molecular Signatures Database Hallmark Gene Set Collection. Cell Syst. 2015;1:417–25.

59. Patro R, Duggal G, Love MI, Irizarry RA, Kingsford C. Salmon provides fast and bias-aware quantification of transcript expression. Nat Methods [Internet]. 2017; Available from: http://www.ncbi.nlm.nih.gov/pubmed/28263959

60. Dobin A, Davis CA, Schlesinger F, Drenkow J, Zaleski C, Jha S, et al. STAR: Ultrafast universal RNA-seq aligner. Bioinformatics. 2013;29:15–21.

